# Fast and accurate differential transcript usage by testing equivalence class counts

**DOI:** 10.1101/501106

**Authors:** Marek Cmero, Nadia M Davidson, Alicia Oshlack

## Abstract

RNA sequencing has enabled high-throughput and fine-grained quantitative analyses of the transcriptome. While differential gene expression is the most widely used application of this technology, RNA-seq data also has the resolution to infer differential transcript usage (DTU), which can elucidate the role of different transcript isoforms between experimental conditions, cell types or tissues. DTU has typically been inferred from exon-count data, which has issues with assigning reads unambiguously to counting bins, and requires alignment of reads to the genome. Recently, approaches have emerged that use transcript quantifications estimates directly for DTU. Transcript counts can be inferred from ‘pseudo’ or lightweight aligners, which are significantly faster than traditional genome alignment. However, recent evaluations show lower sensitivity in DTU analysis. Transcript abundances are estimated from equivalence classes (ECs), which determine the transcripts that any given read is compatible with. Here we propose performing DTU testing directly on equivalence class read counts. We evaluate this approach on simulated human and drosophila data, as well as on a real dataset through subset testing. We find that ECs counts have similar sensitivity and false discovery rates as exon-level counts but can be generated in a fraction of the time through the use of pseudo-aligners. We posit that equivalent class counts is a natural unit on which to perform many types of analysis.

## Introduction

RNA sequencing with short read sequencing technologies (RNA-seq) has been used for over a decade for exploring the transcriptome. While differential gene expression is one of the most widely used applications of this data, significantly higher resolution can be achieved by using the data to explore the multiple transcripts expressed from each gene locus. In particular, it has been shown that each gene can have multiple isoforms, sometimes with distinct functions, and the dominant transcript can be different across samples^1^. Therefore, one important analysis task is to look for differential transcript usage (DTU) between samples.

DTU can be inferred through differential exon usage, where the proportions of RNA-Seq fragments aligning to each exon change relative to each other between biological groups. Anders et al.^2^ showed that exon counts could be used to test for differential exon usage with a generalized linear model that accounts for biological variability. However, counting fragments across exons is not ideal because many fragments will align across multiple exons making their assignment to an individual exon ambiguous. Moreover, individual exons often need to be partitioned into multiple disjoint counting bins when exon lengths differ between transcripts. Typically, there will be more counting bins than transcripts (Figure 2a), resulting in lower power to detect differences between samples.

**Figure 1.**
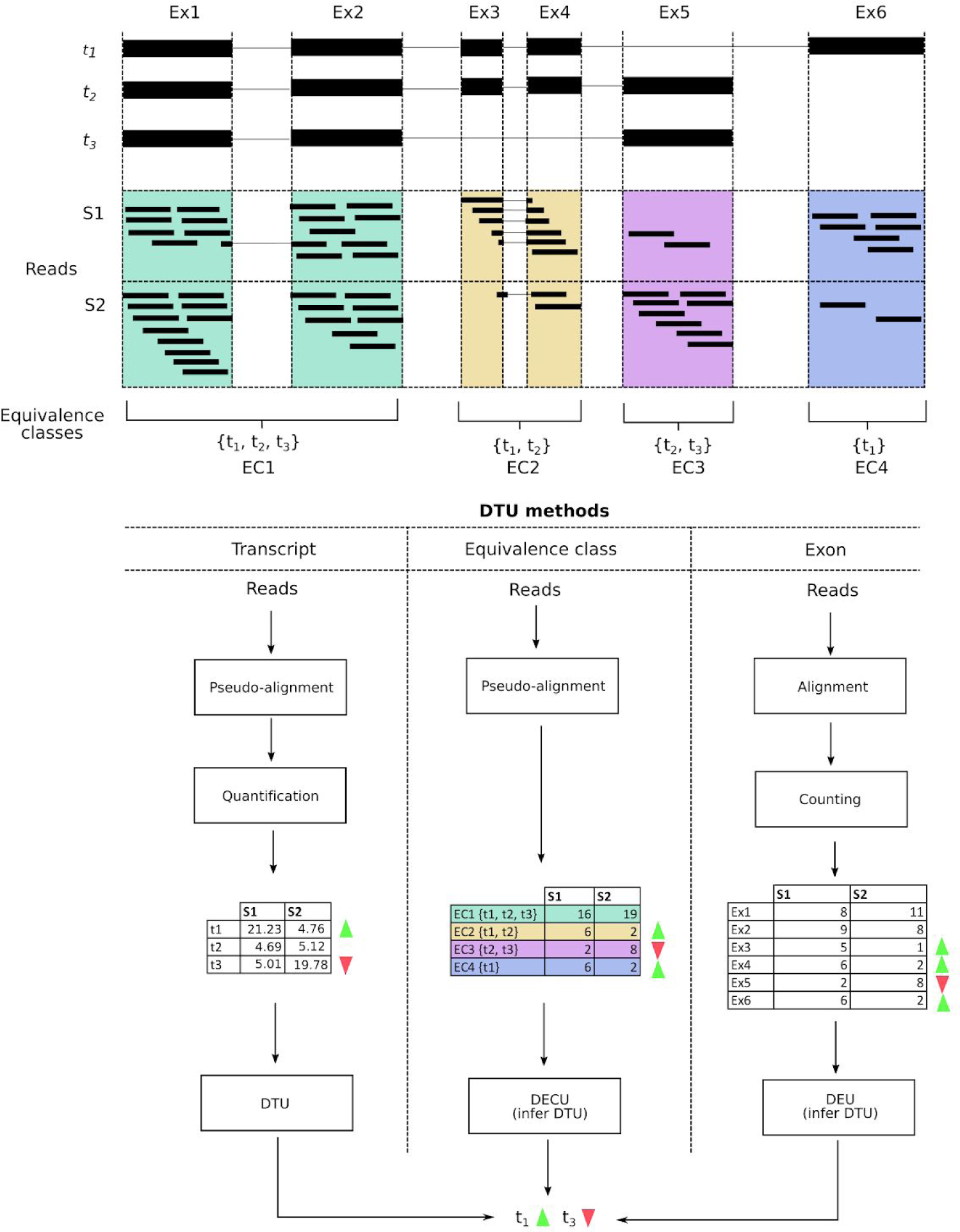
The use of equivalence classes for detecting differential transcript usage (DTU) in a hypothetical gene. The example shows a gene consisting of six exons (Ex1-6) and three transcripts (t_1-3_) resulting in four equivalence classes (EC1-4). t_1_ is predominantly expressed in condition 1 (S1), whereas t_3_ is predominantly expressed in condition 2 (S2). The DTU is evident as a change in the counts for EC2, EC3 and EC4 between conditions. The pipelines for the three alternative methods for detecting DTU are shown: quantification of transcript expression followed by DTU testing, assignment of read counts to equivalence classes followed by testing of equivalent class counts (DECU) and assignment of read counts to exons followed by differential exon counts (DEU). Genes that are detected to have DECU or DEU are inferred to have DTU. The transcript quantification table in the left-most column is example data only, and is not based on real inference.

**Figure 2.**
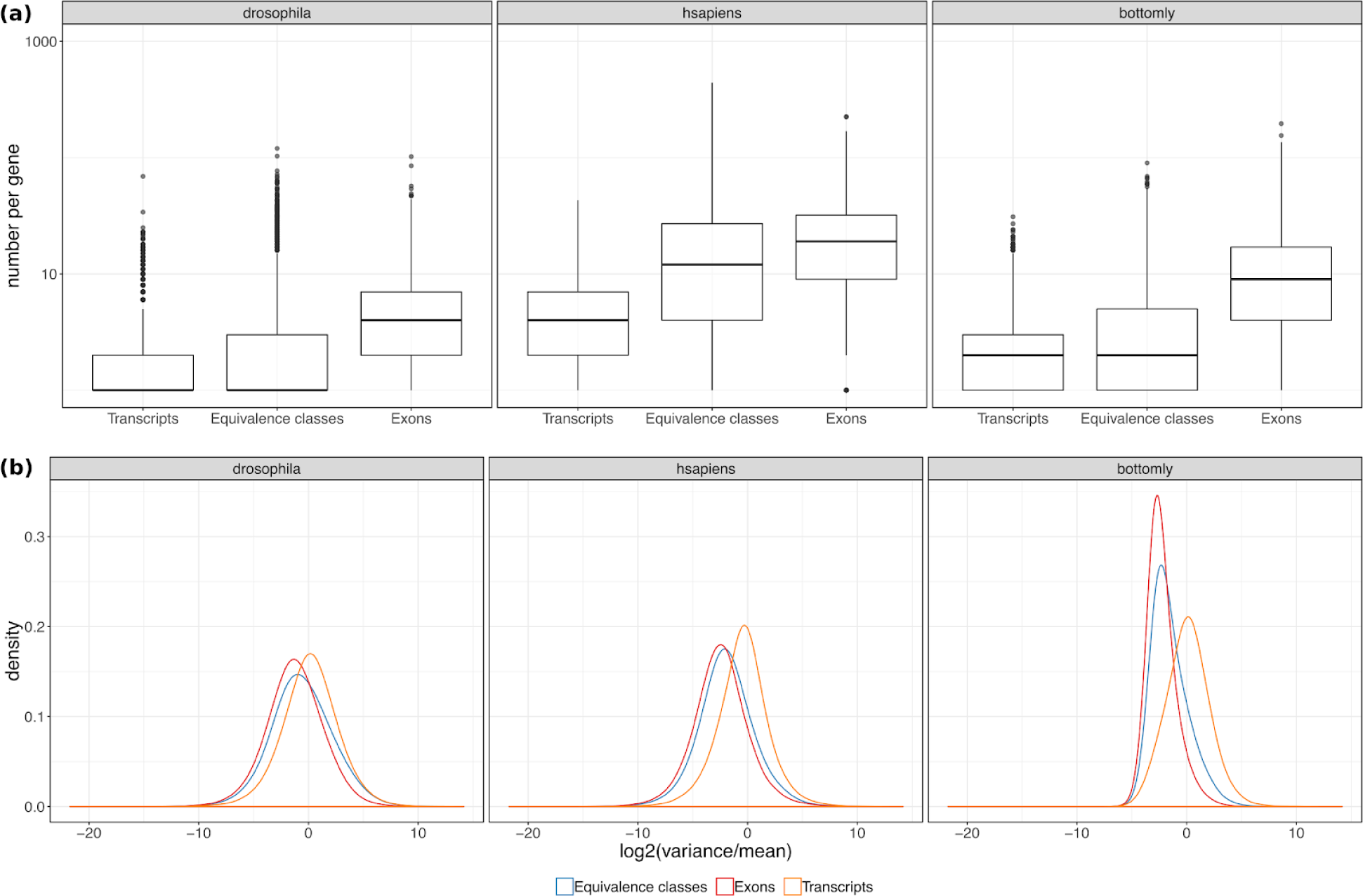
**(a)** The number of transcripts, equivalence classes and exons per gene, where each feature has at least one associated read. **(b)** The density of the log_2_ of the variance of counts over the mean for each feature (calculated per condition).

An alternative to using exon counts for testing DTU is to perform tests directly on estimated transcript abundances^3^. Recently, fast and accurate methods for quantifying gene expression at the transcript level have been developed^4,5^. These methods use transcript annotations that include multiple known transcript sequences for each gene as a reference for the alignment. The lightweight or ‘pseudo’ alignment then assigns each read as ‘compatible’ with one or more transcripts that are a close alignment to the read. Because different transcripts of the same gene share large amounts of sequence, many reads are compatible with several transcripts. A read is then assigned to an equivalence class, which reflects the combination of transcripts compatible with the read sequence (Figure 1). For the purposes of this work, we consider an equivalence class to be defined as in Bray et al.^4^, i.e. any fragments that are pseudo-aligned to the same set of transcripts are considered to be part of the same equivalence class. Figure 1 shows a toy example of a gene with three different transcripts. Depending on its sequence, a read can align to all three transcripts, only two of the transcripts or just one transcript. These different combinations result in four possible equivalent classes, containing read counts, for this gene.

The expression levels of individual transcripts can be estimated from pseudo-aligned reads using all equivalence class counts that are associated with a specific gene^6^. These transcript abundance estimates can be used as an alternative starting measure for DTU testing. It has been shown that estimated transcript abundances can perform well in detecting differential transcript usage^3^, in addition, pseudo-alignment is significantly faster than methods that map to a genome. However, in the most comprehensive comparison using simulated data, exon-count based methods were shown to have slightly better performance^3^.

Here we propose that DTU can be more accurately detected using equivalence class counts directly. Rather than using these counts to first estimate individual transcript abundances before performing DTU, we investigate the potential of performing DTU before the transcript expression estimation step. In this scenario, count-based DTU testing procedures such as DEXSeq are applied directly to alignments generated from fast lightweight aligners, such as Salmon and Kallisto. DTU testing on equivalence classes is fast and alleviates shortcomings in directly estimating transcript abundances before statistical testing. Indeed, performing analysis directly on equivalent classes has been proposed previously in the context of fast clustering single-cell RNA-seq data^7^.

We evaluate the performance of DTU testing on equivalence class counts using real and simulated data, and show that the approach yields higher sensitivity and lower false discovery rates than estimating counts from transcript abundances, and performs as well or better than counting across exons.

## Results

Here we propose an alternative method for performing DTU and evaluate its performance using simulated and real datasets. The method we propose is to first perform alignment with a lightweight aligner and extract equivalence class (EC or transcript compatibility) counts. These ECs are assigned to genes using the annotation of the transcripts matching to the EC. Next, each gene is tested for DTU between conditions using a count based statistical testing method where exon counts are replaced with EC counts (Figure 1). Significant genes can then be interpreted to have a difference between the relative abundance of transcripts of that gene between conditional groups. In evaluating the EC approach, we used Salmon for pseudo-alignment and DEXSeq for differential testing. We then compared DTU results against the alternative quantification and counting approaches, also using DEXSeq for testing (see Methods).

The datasets we used to evaluate performance were simulated data from human and drosophila from Soneson et al.^3^ and biological data from Bottomly et al.^8^. Each of the Soneson datasets consisted of two sample groups, each with three replicates, where 1000 genes were randomly selected to have DTU such that the expression levels of the two most abundant transcripts were switched. The Bottomly dataset contains 10 replicates each from two mouse strains that were used to call truth and then were subsampled to three replicates in the testing scenarios.

### Fewer equivalence classes are expressed than exons

The number of counting bins used for DTU detection has an impact on sensitivity. More bins leads to lower average counts per bin and therefore lower statistical power per bin and more multiple testing correction. We therefore examined the number of ECs, transcripts and exons present in each dataset. Although the theoretical number of ECs from a set of transcripts can be calculated from the annotation and has the potential to be large, not all combinations of transcripts exist or are expressed. The number of equivalence classes calculated from pseudo-alignment depends on the experimental data as only ECs with reads assigned to them are reported. We compared the number of transcripts and exons in the three datasets (with at least one read) to the number of ECs. In both the simulated human and drosophila datasets, as well as in the Bottomly mouse data, the number of ECs is greater than the number of transcripts, but substantially fewer than the number of exons, indicating that there might be more power for testing DTU using ECs, compared to exon counts (Figure 2a).

### Equivalence class replicates have low variance

In addition, we found that the variability of counts across replicates calculated from ECs was lower than that from estimated transcript abundances (Figure 2b), with an average variance to mean ratio of 1.104 in ECs compared to 4.116 in estimated transcript abundance in the Bottomly data. Exon counts had the lowest average variance-mean ratio of 0.428. The simulated human data followed a similar trend, with highest variance to mean ratio for transcripts and the lowest for exons, with ECs displaying a ratio closer to exons than transcripts. In the simulated drosophila data, the variance to mean ratio of ECs was closer to transcripts (with means of 7.652, 6.194 and 2.686 for transcripts, ECs and exons respectively). Supplementary Figure 1 shows the dispersion-mean trends, again demonstrating lower dispersion in ECs compared to transcript abundance estimates. We hypothesise that the greater dispersion observed for transcript data arises from the abundance estimation step used by pseudo-aligners to infer transcript counts. Due to the lower dispersion, we anticipate that ECs yield greater power for DTU compared to transcript abundance estimates.

### Performance of equivalence classes for DTU detection

Several methods were previously tested on the simulated data from Soneson et al.^3^; DEXSeq’s default counting pipeline and featureCounts were shown to perform best. We recalculated exon counts using DEXSeq’s counting pipeline (as recommended by Soneson et al., we excluded region of genes that overlapped on the same strand in the input annotation), and ran Salmon^5^ to obtain both transcript abundance estimates and equivalence class counts. All other comparison results were obtained from Soneson et al.^3^. For the simulated datasets, we found that ECs had the highest sensitivity in both the drosophila and human datasets (Figure 3a) with a TPR of 0.697 and 0.739 respectively (FDR < 0.05). However, ECs also had a slightly higher FDR compared to exon-counting methods.

**Figure 3.**
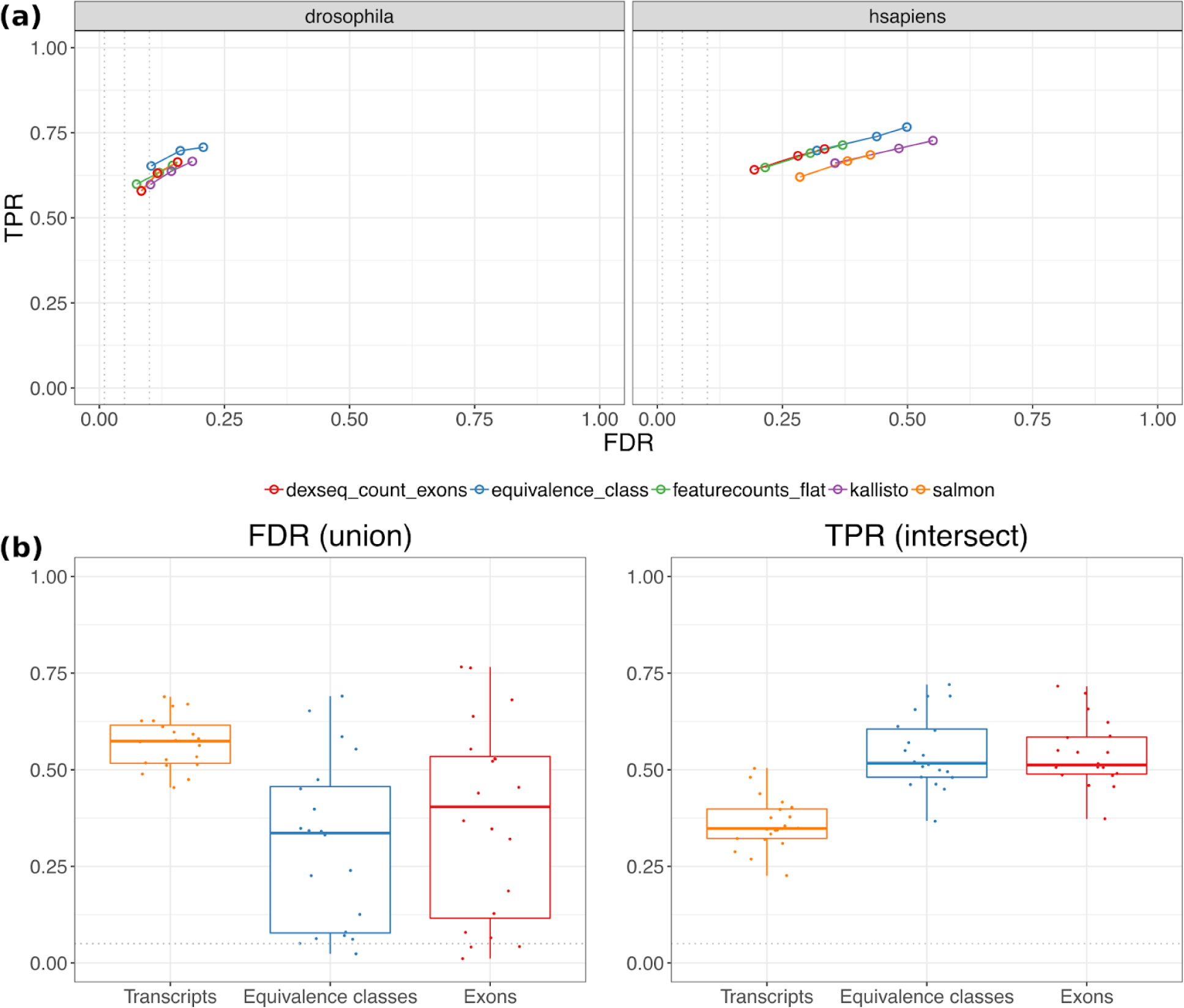
**(a)** The performance of the equivalence class method for differential transcript usage, compared to other state-of-the-art methods on simulated data described in Soneson et al.^3^. **(b)** The ability of the equivalence class, transcript and exon-based methods to recreate the results of a full comparison (10 vs. 10) of the Bottomly data, using only a (randomly selected) subset of samples (3 vs. 3) across 20 iterations. The union of all genes called as significant across all three methods is used to calculate the FDR, and the intersect (genes called by all three methods) is used for the TPR. Full results (union, intersect and each method’s individual truth set) is shown in Supplementary Figure 3.

We next tested the performance of the EC method on a biological dataset from Bottomly et al. We tested the complete RNA-seq dataset (10 vs. 10) for DTU using DEXseq on counts generated from transcript abundance estimates, exons and ECs. To calculate the FDR, we considered the set of ‘true’ DTU genes to be the union of all genes called significant (FDR < 0.05) across the three methods. To calculate the TPR, the intersect of genes called by all methods was used. Supplementary Figure 2 shows the number of significant genes and overlap between all three methods. ECs called the highest number of genes with significant DTU (1485 genes, in contrast to the 748 and 391 genes called significant by the transcript and exon-based methods respectively). Similar to the FDR experiments described in Pimentel et al.^9^, we randomly selected three samples per condition and performed DTU using all three methods and repeated this for 20 iterations. Figure 3b shows the results. EC-based testing performed the best, with a mean FDR of 0.305 across all iterations (compared to a mean FDR of 0.569 and 0.373 for the transcript and exon-based methods respectively). The mean TPR was also slightly higher for ECs at 0.544, compared to exons at 0.539 and 0.36 for the transcript-based method. Results for all three combinations of the ‘truth gene’ sets (union, intersect and individual) are shown in Supplementary Figure 3. The EC-based method had consistently lower FDR, which is also illustrated by the rank-order plot (Supplementary Figure 4), showing the number of false positives present in the top 500 FDR-ranked genes. In terms of the TPR, ECs performed better than transcripts, but worse than exons when using the union of all methods as the truth set. In the Bottomly analysis, Salmon was used as a representative method for transcript abundance estimation. We also performed the analysis with Kallisto, which gave consistent result to Salmon (Supplementary Figure 5).

### Computational performance

While the performance of EC counts in term of sensitivity and FDR are only slightly better than exons level counts, another advantage of using ECs for analysis is the speed of alignment. The process can be broken down into workflow components that include alignment of sequenced reads, quantification and testing. Table 1 shows the compute times for all three methods on all three datasets broken down into workflow components. For the exon counting method, STAR was used for the alignment of reads to the genome (see Methods). In every case, the transcript quantification method was the fastest in terms of total run time followed by ECs and then exons. The difference was mainly driven by the speed of using pseudo alignment for transcript and EC quantification, indicating that for larger datasets the speed of analysis will be significantly faster for our proposed EC based method compared with traditional exon counting methods. A small amount of extra time was also needed for the the EC method for matching EC counts to genes. In addition, DEXSeq generally runs more slowly with larger numbers of counting bins, which is the case for ECs compared with transcripts and improved scalability of DTU approaches is likely to narrow this performance gap. We also note that the transcript-abundance inference stage performed by pseudo-aligners is not necessary for EC-based DTU testing; thus the option to skip this stage would also decrease the compute time.

**Table 1.** Compute times in hh:mm:ss for the simulated data (101 bp paired-end) and Bottomly (76 bp single-end) read data, with each sample aligned and quantified in parallel with access to 256GB RAM and 8 cores per sample, and post-quantification steps performed on count data from all samples from each batch in a single run with 256GB RAM and 8 cores. The drosophila and human samples contained 25M and 40M reads respectively, and the Bottomly sample contained approximately 16M reads.

| **drosophila** | **Transcripts** | **ECs** | **Exons** |
| --- | --- | --- | --- |
| Alignment | - | - | 0:29:26 |
| Quantification | 0:06:44 | 0:06:44 | 0:28:21 |
| Match ECs | - | 0:00:16 | - |
| DEXSeq DTU | 0:01:06 | 0:02:25 | 0:03:20 |
| **Total** | **0:07:50** | **0:09:25** | **1:01:07** |

| **hsapiens** | **Transcripts** | **ECs** | **Exons** |
| --- | --- | --- | --- |
| Alignment | - | - | 0:10:54 |
| Quantification | 0:04:59 | 0:04:59 | 0:52:13 |
| Match ECs | - | 0:01:26 | - |
| DEXSeq DTU | 0:04:34 | 0:16:22 | 0:15:50 |
| **Total** | **0:09:33** | **0:22:47** | **1:18:57** |

| **mouse (Bottomly)** | **Transcripts** | **ECs** | **Exons** |
| --- | --- | --- | --- |
| Alignment | - | - | 0:01:24 |
| Quantification | 0:00:32 | 0:00:32 | 0:07:44 |
| Match ECs | - | 0:01:34 | - |
| DEXSeq DTU | 0:07:40 | 0:14:28 | 0:34:00 |
| **Total** | **0:08:12** | **0:16:34** | **0:43:08** |

We also considered peak RAM usage (shown in Supplementary Table 1), and alignment was found to use the most RAM. Overall, methods utilising pseudo alignment required significantly lower memory compared with traditional alignment. For the most RAM intensive dataset, the human simulation, exon counting required 29GB compared to 10GB for ECs and 5GB for estimated transcript abundances.

## Discussion

DTU detection has previously been approached by either testing for changes to the read coverage across exons or changes in the relative abundance of transcripts. These approaches are intuitive but are not necessarily optimal for short read data analysis. In particular, individual exons are not necessarily the optimal unit of isoform quantification as there are often many more exons than transcripts. In addition, transcript quantification can be difficult because read assignment is ambiguous. Fortunately, transcript quantification methods generate equivalence class counts as a forestep to estimating abundances. We propose that equivalence classes are the optimal unit for performing count based differential testing. Equivalence class counts benefit from the advantages of both exon and transcript counts: they can be generated quickly through pseudo-alignment, there are fewer expressed than exons, and they retain the low variance between replicates seen in exon counts compared to transcripts abundances.

Here we evaluated the use of equivalent classes as the counting unit for differential transcript usage. We used two simulated datasets from drosophila and human and one biological dataset from mouse. Our results suggest that equivalent class counts provide equal or better accuracy in DTU detection compared to exon counts or estimated transcript abundances. We also found the analysis was quick to run and we provide code to convert pseudo alignments into gene level EC annotations.

The ECs used in our evaluation are defined using only the set of transcripts for which reads are compatible. Extensions to this model have been proposed that incorporate read-level information, such as fragment length, to more accurately calculate the probability of a read arising from a given transcript^10^. Although, we do not consider probability-based equivalence classes in this work, incorporating this information for DTU deserves exploration in future work. In addition, EC counts may be calculated from full read alignment rather than pseudo-alignment^11,12^, which has the potential to improve accuracy further.

One limitation of using equivalence classes, however, is in the interpretation of the results. Although we can detect DTU at the gene-level, it is not simple to determine which isoforms have changed abundance without further work. We propose that superTranscripts^13^, which are a method for visualising the transcriptome, could be used for interpretation. Alternatively, transcript abundances, which are generated together with ECs, can still be used to provide insight into the isoform switching.

Finally, in this work, we have focused on differential transcript usage, but EC counts have the potential to be useful in a range of other expression analysis. In particular, EC counts could be used as the initial unit of measurement for many other types of analysis such as dimension reduction visualisations, clustering and differential expression.

## Methods

### Obtaining count data

We detected sequence content bias in the Bottomly RNA-seq data using FastQC v0.11.4, and therefore performed trimming using Trimmomatic^14^ 0.35, using recommended parameters (http://www.usadellab.org/cms/?page=trimmomatic). The simulated Soneson data was not trimmed.

To obtain EC and transcript abundance counts, Salmon^5^ v0.12.0 was run on the drosophila, human and Bottomly datasets in quant mode using default parameters and the *--dumpEq* argument to return EC output. Kallisto^4^ was run in *pseudo* mode with the *--batch* argument to run all samples simultaneously. Fragment length and standard deviation were estimated from all reads of a single sample from the Bottomly data (SRR099223). Equivalence classes were then matched between samples and compiled into a matrix using the python scripts (create_salmon_ec_count_matrix.py and create_kallisto_ec_count_matrix.py), available in the paper github repository (below).

To perform the exon-based counts, raw reads were first aligned using STAR^15^ v2.5.2a, then the DEXSeq-count annotation was prepared excluding overlapping exon-parts, from different genes, on the same strand (--aggregate=‘no’). DEXSeq-count was then run using default parameters. The same genome and transcriptome references for drosophila and human were used as in Soneson et al.^3^, with only protein-coding transcripts considered for the Salmon index. For the Bottomly data, we used the NCBIM37 mm9 mouse genome and Ensembl release 67 transcriptome. Non-protein-coding transcripts were filtered out, as with the Soneson transcriptome reference.

### DTU analyses

All the code to run the analyses and generate the paper figures from the count matrices is available at https://github.com/Oshlack/ec-dtu-paper. Equivalence classes mapping to more than a single gene were removed. No other filtering was performed on any of the data types. DEXSeq v1.26 was used to run all DTU analyses.

### Datasets

The Soneson et al.^3^ drosophila and human simulation data was obtained from ArrayExpress repository with accession number E-MTAB-3766 (http://www.ebi.ac.uk/arrayexpress/experiments/E-MTAB-3766/). Truth data was obtained from: http://imlspenticton.uzh.ch/robinson_lab/splicing_comparison/. The Bottomly et al.^8^ dataset was obtained from the NCBI Sequence Read Archive with accession number of SRP004777 (https://trace.ncbi.nlm.nih.gov/Traces/sra/?study=SRP004777).

