## Supplementary figures and tables for "Fast and accurate differential transcript usage by testing equivalence class counts"

### Supplementary Material

Marek Cmero[1], Nadia Davidson[1*], Alicia Oshlack[1,2*]

1. Murdoch Children's Research Institute, Flemington Road, Parkville, Australia
2. School of Biosciences, Faculty of Science, University of Melbourne, Melbourne, Victoria, Australia

*These authors contributed equally in supervision of this work

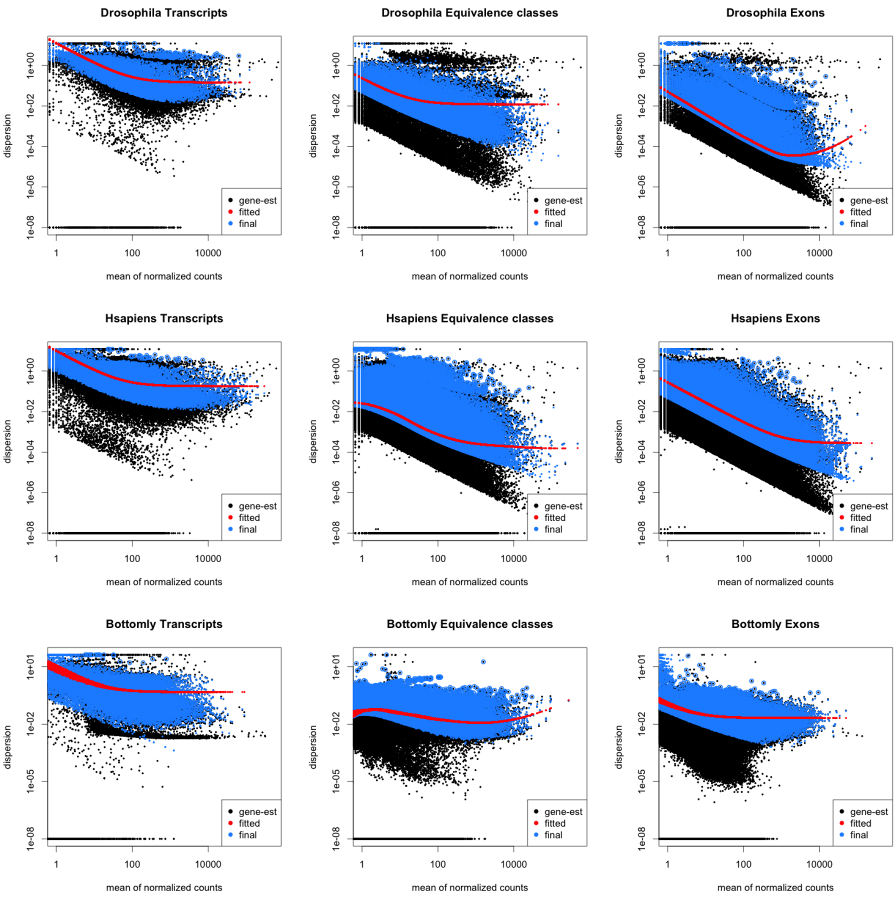

Supplementary Figure 1: Shows the dispersion versus mean normalised counts for all features across the three data sets, generated using DEXSeq's 'plotDispEsts' function. As described in Love et al. [1], the red line shows the fitted dispersion-mean trend, the blue dots indicate the shrunken dispersion estimates, and the blue circles indicate outliers not shrunk towards the prior.

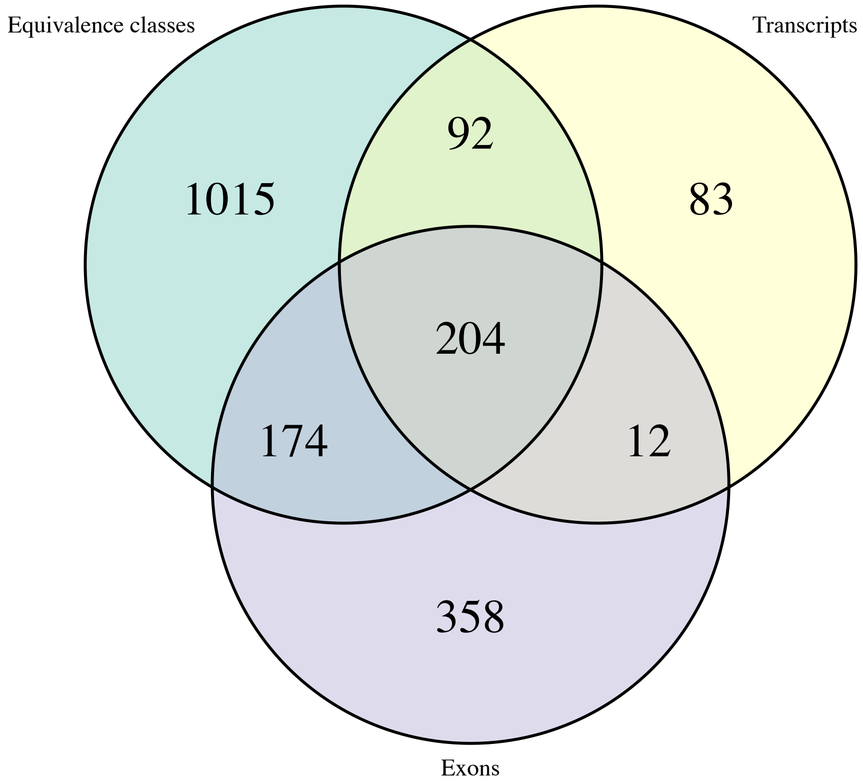

Supplementary Figure 2: Shows the significant genes (FDR $< 0.05$) shared between the methods, obtained from DEXSeq run on the full Bottomly et al. [2] data set for each feature.

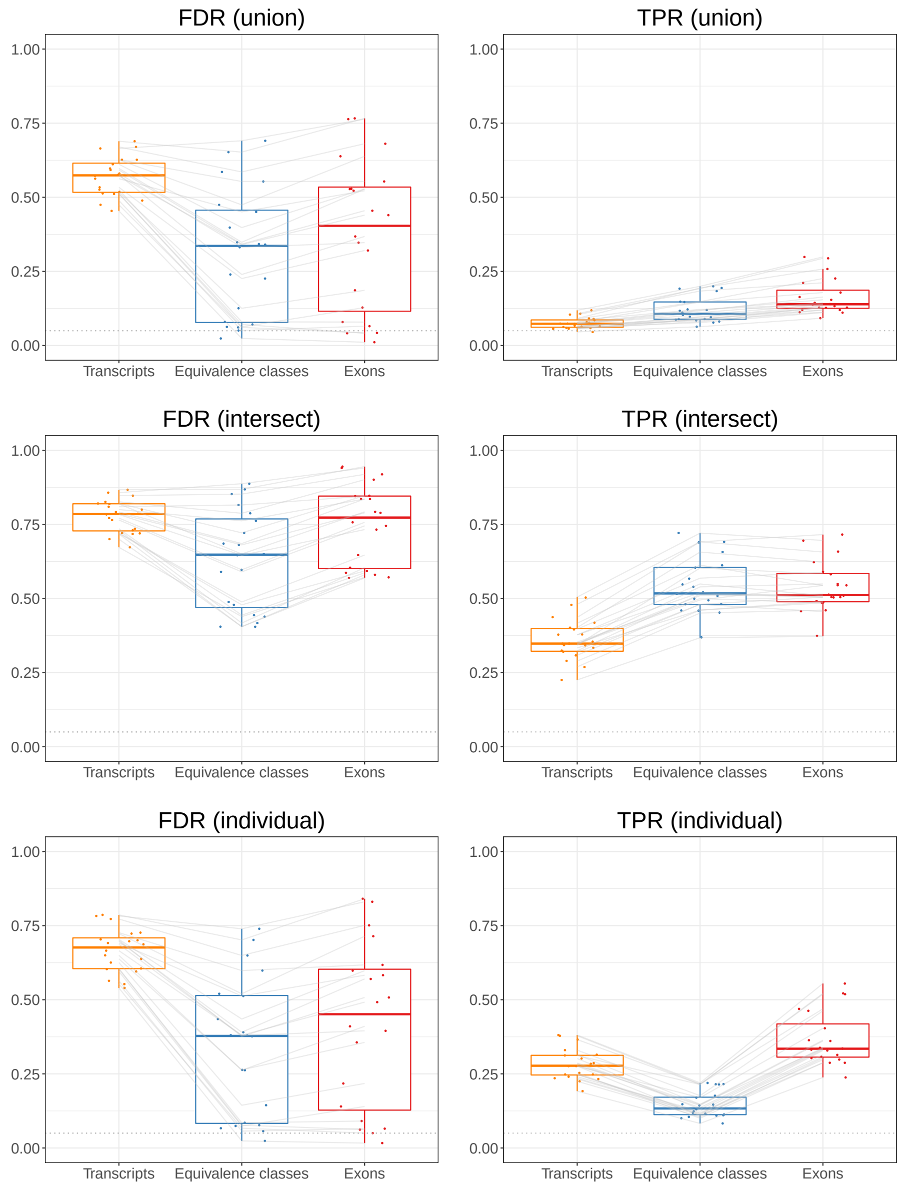

Supplementary Figure 3: Shows the ability of the three methods to recreate the results of a full comparison (10 vs. 10) of the Bottomly et al. [2] data using random subsets of 3 vs. 3 samples across 20 iterations. The lines between the plots join data points from the same iteration. Each row uses a different 'truth' set: union is the set of genes called significant by any method, intersect is the set of genes called significant by all methods, and individual is the set of genes called significant by that method only.

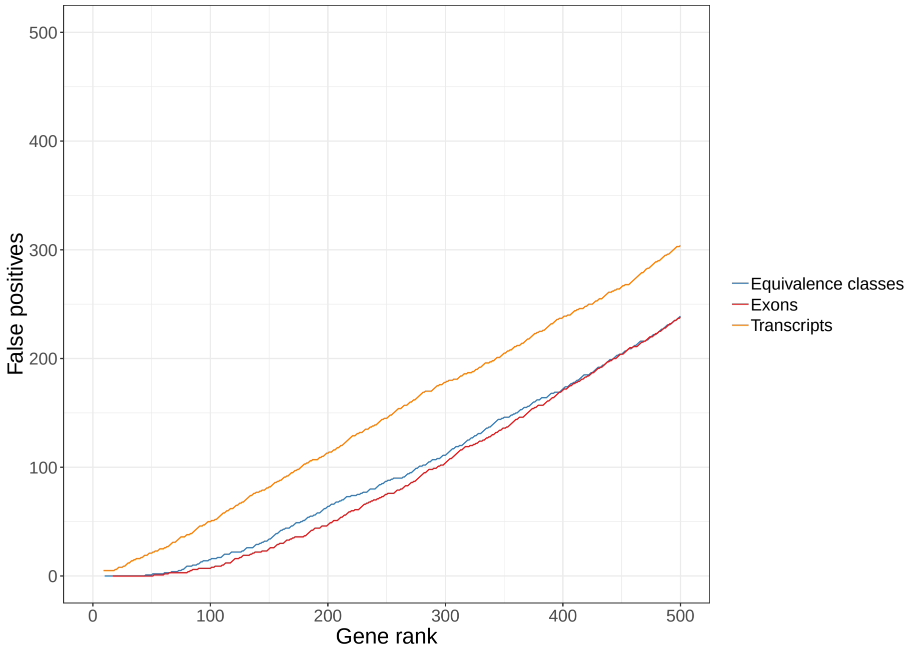

Supplementary Figure 4: The number of false positives versus each gene's rank (by FDR) for one iteration (3 vs. 3) of the Bottomly subset tests for the top 500 genes. The union of significant genes across all methods was used as the truth set.

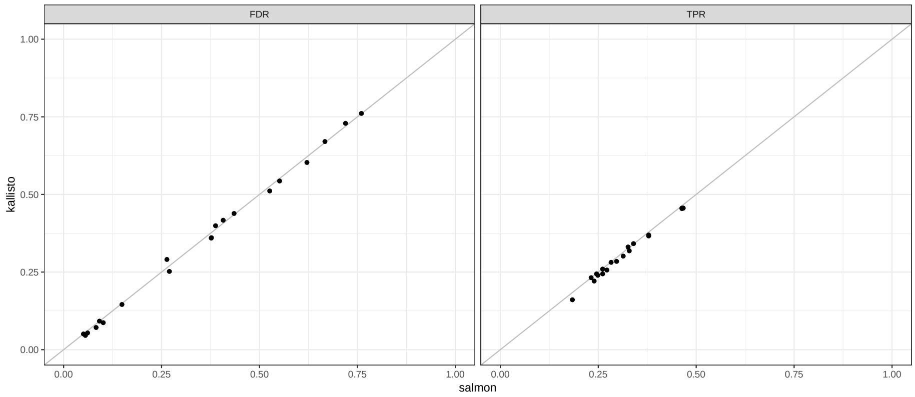

Supplementary Figure 5: Kallisto [3] versus salmon's [4] performance on the Bottomly subset testing experiments, using each method's significant genes from the full (10 vs. 10) run as the truth set for calculating both metrics.

Supplementary Table 1: Maximum RAM usage for each job in GB. Each task was run as specified in the compute times table in the main paper (Table 1).

| **drosophila** | **Transcripts** | **ECs** | **Exons** |
|---|---|---|---|
| Alignment | - | - | 6.55 |
| Quantification | 1.87 | 1.87 | 0.50 |
| Match ECs | - | 0.53 | - |
| DEXSeq DTU | 1.22 | 1.22 | 1.39 |
| **Max** | 1.87 | 1.87 | 6.55 |

| **hsapiens** | **Transcripts** | **ECs** | **Exons** |
|---|---|---|---|
| Alignment | - | - | 28.54 |
| Quantification | 5.15 | 5.15 | 1.11 |
| Match ECs | - | 4.02 | - |
| DEXSeq DTU | 2.19 | 9.95 | 4.84 |
| **Max** | 5.15 | 9.95 | 28.54 |

| **mouse (Bottomly)** | **Transcripts** | **ECs** | **Exons** |
|---|---|---|---|
| Alignment | - | - | 24.73 |
| Quantification | 2.54 | 2.54 | 0.35 |
| Match ECs | - | 2.38 | - |
| DEXSeq DTU | 2.15 | 7.10 | 5.48 |
| **Max** | 2.54 | 7.10 | 24.73 |
